# Torsional Twist of the SARS-CoV and SARS-CoV-2 SUD-N and SUD-M domains

**DOI:** 10.1101/2024.08.13.607777

**Authors:** Monica Rosas-Lemus, George Minasov, Joseph S. Brunzelle, Taha Y. Taha, Sofia Lemak, Shaohui Yin, Ludmilla Shuvalova, Julia Rosecrans, Kanika Khanna, H Steven Seifert, Alexei Savchenko, Peter J. Stogios, Melanie Ott, Karla J. F. Satchell

## Abstract

Coronavirus non-structural protein 3 (nsp3) forms hexameric crowns of pores in the double membrane vacuole that houses the replication-transcription complex. Nsp3 in SARS-like viruses has three unique domains absent in other coronavirus nsp3 proteins. Two of these, SUD-N (Macrodomain 2) and SUD-M (Macrodomain 3), form two lobes connected by a peptide linker and an interdomain disulfide bridge. We resolve the first complete x-ray structure of SARS-CoV SUD-N/M as well as a mutant variant of SARS-CoV-2 SUD-N/M modified to restore cysteines for interdomain disulfide bond naturally lost by evolution. Comparative analysis of all structures revealed SUD-N and SUD-M are not rigidly associated, but rather, have significant rotational flexibility. Phylogenetic analysis supports that the disulfide bond cysteines are also absent in pangolin-SARS and closely related viruses, consistent with pangolins being the presumed intermediate host in the emergence of SARS-CoV-2. The absence of these cysteines does not impact viral replication or protein translation.

## INTRODUCTION

The coronaviruses SARS-CoV and SARS-CoV-2 are the causative agents of Severe Acute Respiratory Syndrome (SARS) and Coronavirus disease 2019 (COVID-19), respectively [1-3]. Both viruses caused global outbreaks in humans in the early 21^st^ century, with SARS accounting for more than 8000 probable cases and 774 deaths from 2002-2004 in Asia and North America and COVID-19 accounting for over 7 million deaths worldwide from 2019-2024 [4, 5]. Coronavirus are enveloped, positive-strand, single-stranded RNA (ssRNA+) viruses. Upon delivery of the ssRNA into infected cells, non-structural proteins (nsp) translated by host ribosomes from the incoming RNA strand are produced, leading to modification of the endoplasmic reticulum to create double membrane vesicles (DMVs) that ultimately contains the viral-encoded replication-transcription complex (RTC). Within the DMVs, the RTC generates genomic ssRNA+ copies for packaging into new viral capsids along with subgenomic messenger RNA (mRNA) strands that will be translated by host ribosomes to produce the viral structural S, N, E, and M proteins [6]. Although the RNA molecules are produced within the DMV, both packaging of genomic RNA into capsids and translation of subgenomic RNAs requires the RNA molecules be exported from the DMV to the cytoplasm [6-8]. Cryo-electron tomography of DMV structures has revealed that the DMV membranes contain hexameric transmembrane pores formed by non-structural protein 3 (nsp3) in complex with non-structural protein 4 (nsp4) and non-structural protein 6 (nsp6) and it is postulated that viral RNAs pass through these pores to access the cytoplasm for translation or viral capsid packaging [9, 10].

Nsp3 is the largest protein produced by coronaviruses [11, 12]. The hexamer of nsp3 extends from the DMV into the cell cytoplasm with each nsp3 protein tethered to the DMV by two transmembrane domains, TM1 and TM2. The small nsp3 ectodomain (3Ecto) between TM1 and TM2 is displayed on the luminal side of the DMV and interacts with nsp4 as part of the transmembrane pore complex [9-12]. The C-terminal Y1 and CoV-Y1 domains are also exposed to the cytoplasmic side of the DMV (Figure 1A).

**FIGURE 1.**
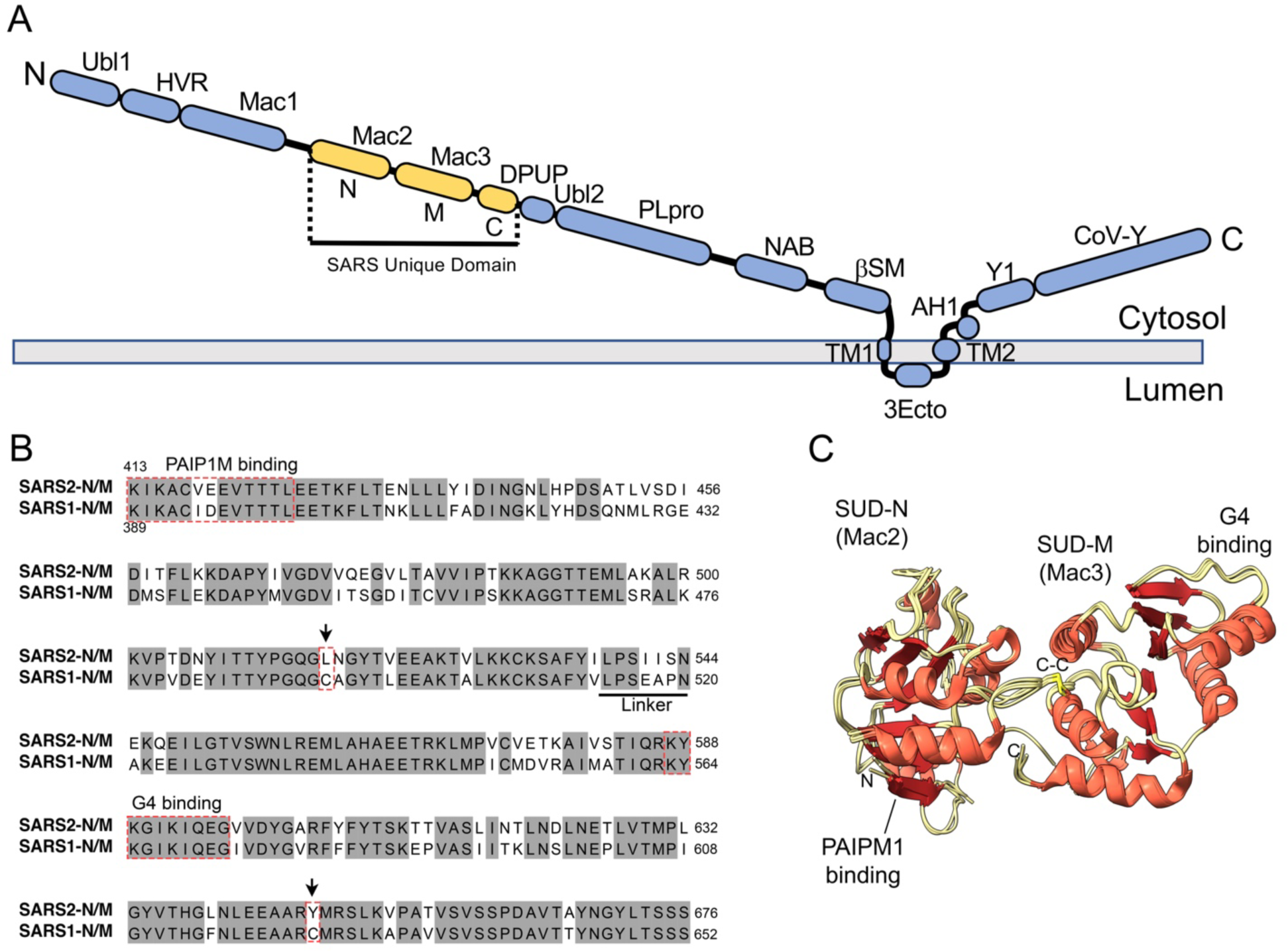
The nsp3 SARS Unique domains from SARS-CoV and SARS-CoV-2. (A) Diagram of the domain structure of nsp3 form SARS-CoV, SARS-CoV-2 and related viruses. Domains in the long N-terminal extension into the cytosol are the ubiquitin-like domain 1 (Ubl1), the hypervariable region (HVR), macrodomain I (Mac1), macrodomain II (Mac2 / SUD-N), macrodomain III (Mac3 / SUD-M), domain preceding Ubl2 and PLpro (DPUP / SUD-C), ubiquitin-like domain 2 (Ubl2), papain-like protease (PLpro), nucleic acid binding (NAB) betacoronavirus-specific marker (βSM). The transmembrane region 1 (TM1) and region 2 (TM2) cross the DMV double membrane with the nsp3 ectodomain (3Ecto) facing the luminal side of the membrane and binds with nsp4. The C-terminal domains Y1 and coronavirus Y (CoV-Y) are exposed to the cytosolic face of the DMV. The SARS Unique domain (SUD) is shown in yellow with N-terminal (N), middle (M), and C-terminal (C) domains marked. (B) Amino acid alignment of SUD-N/M from SARS-CoV and SARS-CoV-2 with important regions that bind PAIP1M and G4 motifs indicated with red dashed boxes. Cysteine residues that participate in disulfide bond in SARS1-N/M substitutions are marked with arrows. (C) Overlay of all six chains of prior determined structures of SARS-CoV SUD-N/M (SARS1-N/M, PDB ID 2w2g and 2wct). Backbone in beige, helices in orange, beta-strands in red. Structures have conserved fold with r.m.s.d. of ∼0.7. All show two macrodomains with six beta-strand wrapped by five helices. Linker is disordered in all structures. Disulfide bond linkage between Cy492 and Cys623 (shown in yellow in center of structures) is seen in all six chains.

The organization of N-terminal domains of nsp3 are variable across different coronavirus but are conserved between SARS-CoV and SARS-CoV-2 (Figure 1A). Three of these domains, known as the SARS unique domains (SUD), are found within nsp3 only in SARS-CoV, SARS-CoV-2 and other closely related SARS-like coronaviruses [11, 13-15], and are notably absent in other human coronaviruses including MERS-CoV, HCoV-OC43, and HCoV-229E [11, 12, 15]. These unique domains include macrodomain 2 (Mac2, also known as SUD-N), macrodomain 3 (Mac3, also known as SUD-M), and the domain-preceding Ubl2 and PLpro (DPUP, also known as SUD-C). The combination of Mac2 and Mac3 (also known as SUD2_Core_) will be referred to here as SUD-N/M and as SARS1-N/M for SARS-CoV and SARS2-N/M for SARS-CoV-2.

The SARS1-N and SARS1-M domains of SARS-CoV each have a macrodomain fold similar to the ADP-ribosylhydrolase domain Mac1 [16-18]. However, unlike Mac1, these domains do not bind ADP-ribose [16]. Rather, the SARS1-N/M proteins bind to RNA and DNA oligonucleotides, especially those folded to form guanine-quadruplex (G4) structures [12, 16]. These three-dimensional structures are formed in guanine-rich regions of nucleotides in which four guanine bases organize into a guanine tetrad plane and then two or more tetrad planes stack to form the G4 [19]. Modeling and site-directed mutagenesis studies of SARS1-N/M suggest both RNA and DNA G4 nucleotide structures bind via lysines in a flexible surface exposed loop of the SARS1-M domain with minor contributions from the SARS1-N domain [20].

The SUD-N/M domains from SARS-CoV are 74% identical and 85% similar in amino acid sequence with the SARS-CoV-2 domains (Figure 1B), and likewise, SARS2-N/M domains bind to G4 nucleotide structures [21, 22]. SARS2-N/M binds preferentially to a G4 motif found in the 5’UTR of *TRF2* mRNA folded to form a G4 structure and this correlates with inhibition of the unfolded protein response that detects endoplasmic reticulum stress [22]. SARS2-N/M also binds to G4 motifs formed by folding of single-stranded DNA, including a G4 motif of *BCL2* and this interaction is suggested to promote apoptosis [22]. Further, SARS2-N/M has been shown to bind the 5’UTR for SARS-CoV-2 RNAs and to possess global RNA binding activity [23].

Interactions of SARS2-N/M domains with other proteins may also have functionally significant relevance to viral infection. The binding of SARS2-N/M domains to G4 structures is enhanced by its interaction with viral non-structural protein 5 (nsp5) and this interaction is also associated with enhanced apoptosis [24]. In addition, both SARS1-N and SARS2-N bind the middle domain of polyadenylate binding protein-interacting protein 1 (PAIP1M), a cytoplasmic co-factor that binds to polyadenylate binding protein 1 (PAB1), which is an enhancer of ribosome translation [21, 25]. The interaction with PAIP1M as well as the co-sedimentation of nsp3 with ribosomal fractions and with ribosomal inhibitor non-structural protein 1 (nsp1) suggest that SUD-N domains may function in part to promote translation of viral mRNAs over host mRNAs [25].

X-ray structures of SARS1-N/M have been determined and reveal these two domains form a bi-lobal protein connected by both a flexible linker peptide (underlined in Figure 1B) and by an interdomain disulfide bridge (PDB ID 2w2g and 2wct) [16]. In this study, we determine additional structures of the SARS1-N/M domains as well as a variant of SARS2-N/M that forms the interdomain disulfide bond. Analysis of all available structures reveal these two domains are more flexible than suggested by earlier structural studies and the domains have torsional twist with respect to each other, although each of the macrodomains are structurally conserved. The presence or absence of cysteines has a minimal contribution to binding to nucleic acids and did not impact viral replication. Across all the SARS-like viruses, the capacity to form the disulfide bond is shared among viruses closely related to SARS-CoV while viruses related to Pangolin-SARS lack the cysteines, consistent with a model of Pangolin-SARS being a progenitor strain for SARS-CoV-2.

## RESULTS

### A complete structure of SARS-CoV SUD-N/M

The x-ray structure of the 259 a.a. joined SARS1-N/M domains was previously solved at 2.22 Å with two chains in the asymmetric unit (PDB ID 2w2g) [16]. A second lower-resolution structure has four chains in the asymmetric unit (PDB ID 2wct) [16]. All six chains from both structures have nearly identical backbone folds (Figure 1C). Since all the prior SARS1-N/M structures are nearly identical (root-mean-squared-deviation (r.m.s.d.=0.6-0.8 Å), only the structure chain reported as PDB ID 2w2g/A will be analyzed in this study.

As part of a structural genomics project to address the COVID-19 pandemic, we attempted to crystallize the recombinant SARS2-N/M domains of the comparable 259 a.a. protein as used for prior structures of protein from SARS1-N/M (Figure 1C). After multiple failed attempts to crystallize SARS2-N/M, we cloned and purified SARS1-N/M to test the compatibility of our approach for protein purification and crystallization using this protein known to crystallize. This SARS-CoV derived protein rapidly crystallized, data were collected, and the structure was solved by molecular replacement using PDB ID 2w2g as a model.

This new structure (PDB ID 8ufl) was determined at 2.51 Å and included two chains in the asymmetric unit, designated Chain 8ufl/A and 8ufl/B. Similar to the prior structures, the SARS1-N has a macrodomain fold comprised of a six stranded β sheet (βN1-βN6-βN5-βN2-βN4-βN3) surrounded by six helices (αN1-αN6) (Figure 2A). Unique to this structure, chain 8ufl/A was complete across all residues and included resolution of residues 515-523 in the peptide linker between the N and M domains, which was disordered in chain B and in all prior structures. The initial strand of the SARS1-M domain in this new structure resolved as a helix missing from all previous structures and here designated as αM1 followed by βM1. This M domain now has a similar macrodomain as the SARS1-N domain with a composition as a six-strand β sheet (βM1-βM6-βM5-βM2-βM4-βM3) surrounded by six helices (αM1-αM6). Similar to all prior structures, this new structure resolved with an interdomain disulfide bond between residues Cys492 linking the large loopN8 between βN5 and αN6 with Cys623 in αM5 of SARS1-M (Figure 2A).

**FIGURE 2.**
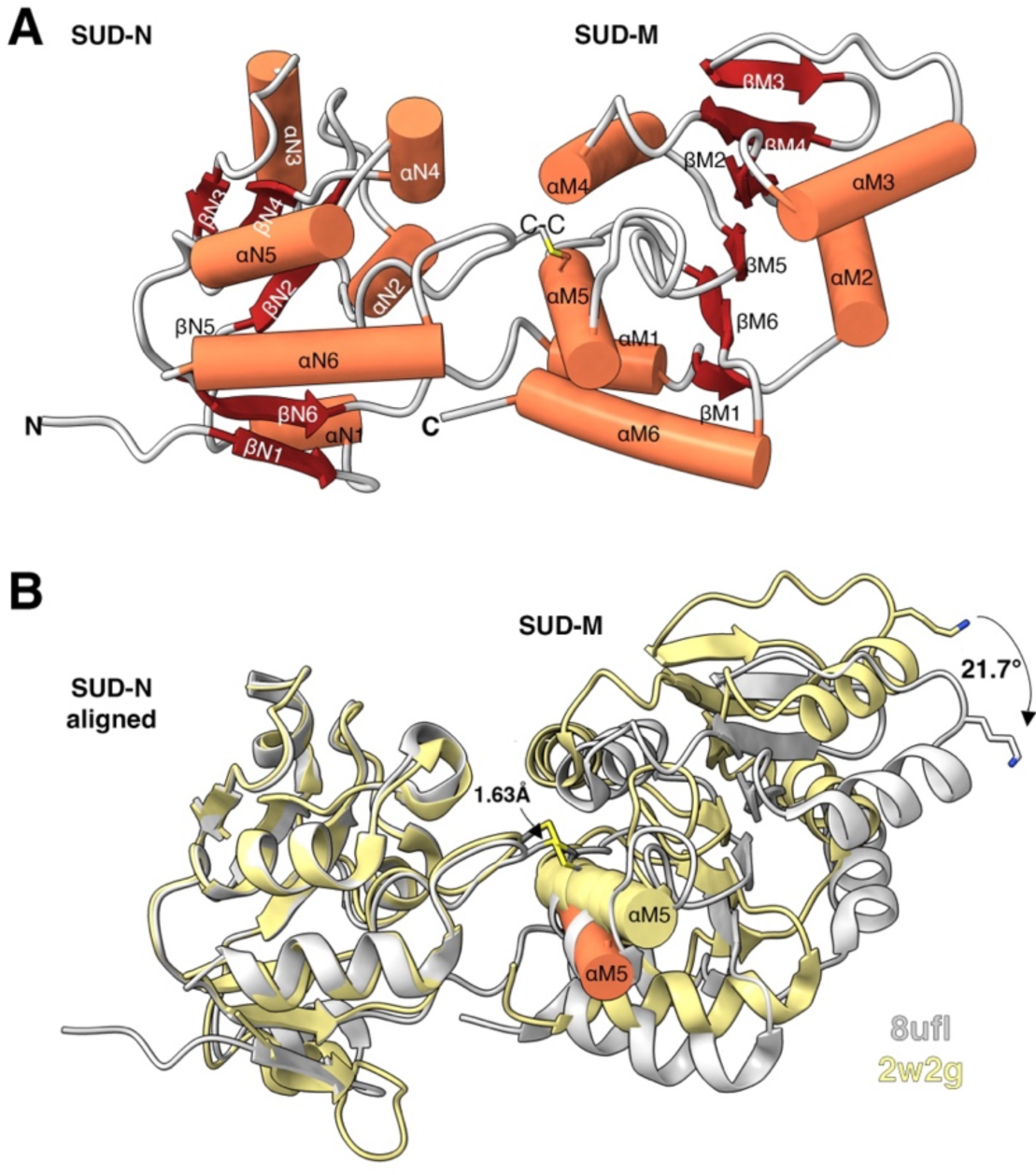
Complete structure of SARS1-N/M. (A) Cylinder and slabs diagram of SARS1-N/M (PDB ID 8ufl/A) with backbone in light grey, helices in coral, and beta-strands in red. Domains are labeled according to this complete structure. Disulfide bond (C-C) between SARS1-N Cys492 and SARS1-M Cys623 is shown (B) Ribbon diagram of SARS1-N/M structure 8ufl (light grey) overlaid with structure 2w2g/A (yellow) by alignment of SUD-N domains. Distance of shift of C-C bond is marked. Helices αM5 are shown in yellow (2w2g) and coral (8ufl) to emphasize degree of shift of this C-C linked helix. Angle of rotation of SUD-M compared to SUD-N is shown at right as measured from Cys492 to Lys545 for each structure.

The other significant difference in this structure compared to prior SARS-CoV structures (Figure 1C) is that the relative rotational orientation of SUD-N and SUD-M was shifted compared to the alignment of all other structures. Although SUD-N and SUD-M each independently aligned between 2w2g and 8ufl, when SUD-N was aligned, SUD-M was rotated 21.7° from reorientation of the disulfide bond and flexibility in the linker. This shift moved Lys565 that participates in binding to G4 quadruplex structures by 12.9 Å.

These data indicated that in contrast to prior studies, there is rotational flexibility in the disulfide bound structures such that the cleft between SUD-N and SUD-M is not a fixed interface and flexibility is introduced by both the disulfide bond and the interdomain peptide linker.

### Structure of SARS-CoV-2 SUD-N/M variant with reintroduced disulfide bond

The successful new structure of SARS1-N/M (Figure 2A), and our high throughput crystallization approaches supported that the failure of SARS2-N/M to crystallize was most likely linked to the 25% difference in amino acid sequence between the two proteins. Analysis of the aligned sequences of these domains revealed that SARS2-N/M lacks the two cysteines that form the interdomain disulfide bond in SARS1-N/M, thus the joined domains could not form this potentially stabilizing disulfide bond (Figure 1B). A new synthetic clone of SARS2-N/M was generated that exchanged codons for residues Leu516 in SARS2-N and Tyr647 in SARS2-M to cysteines to introduce the potential for disulfide bond formation at the same location as found in SARS1-N/M. The protein variant, here referred to as SARS2-N^C^/M^C^, was purified in the absence of reducing agent and set up for crystallization. The protein rapidly crystallized, data were collected, and the structure determined at 1.65 Å resolution. The structure was comprised of one chain in the asymmetric unit.

The SARS2-N^C^ domain resolved a similar macrodomain fold as for SARS1-N (r.m.s.d.=0.857), except that the large loop with αN4 in SARS1-N was disordered (Figure 3A). In the SARS2-N^C^/M^C^ structure, the interdomain linker (residues 541-548) was not resolved and the first strand of the SARS2-M^C^ domain did not contain either the αM1 or βM1 that were determined in the complete SARS1-N/M structure. The SARS2-M domain is otherwise structurally conserved with SARS1-M (r.m.s.d.=0.617).

**FIGURE 3.**
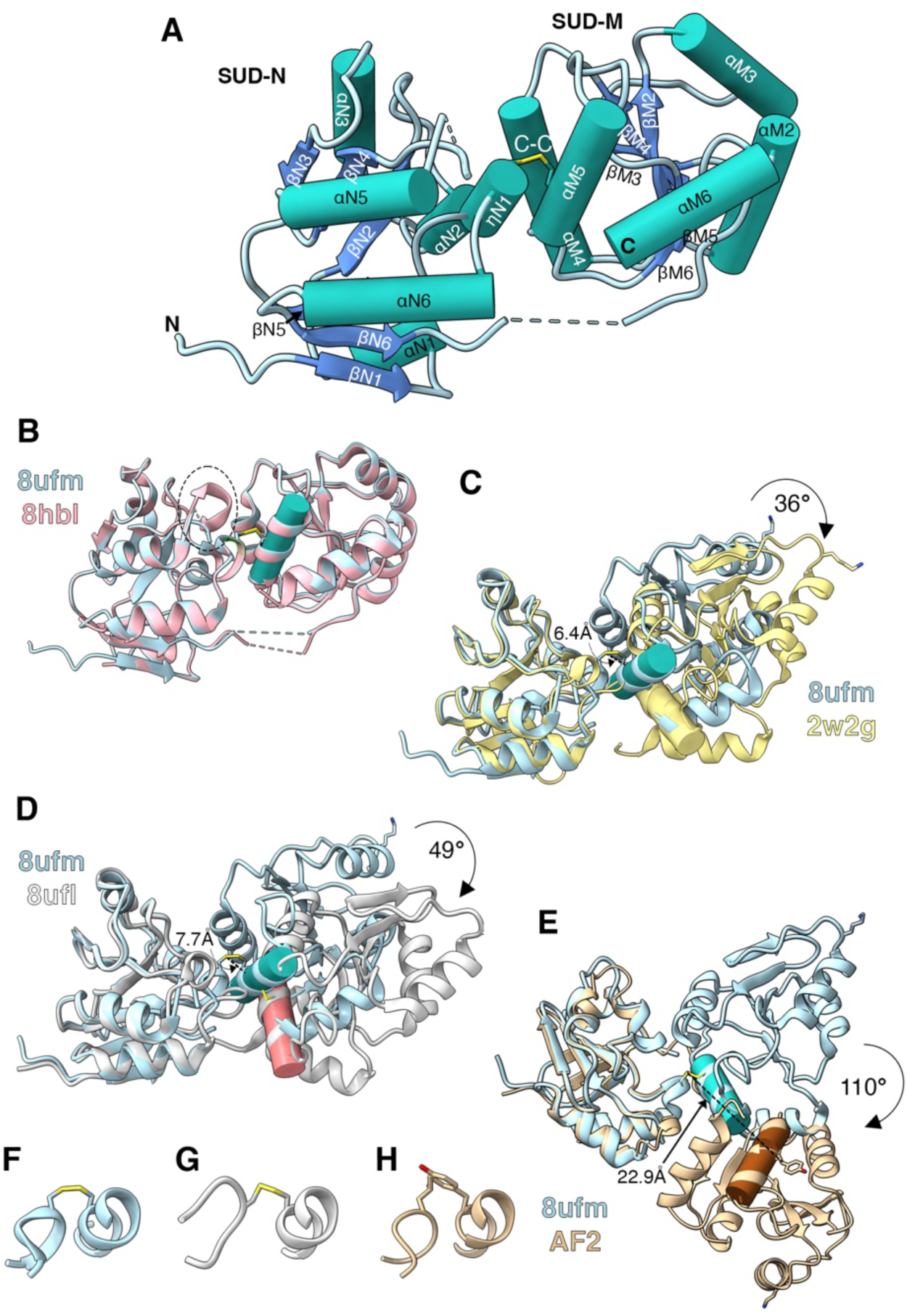
Structure SARS2-N^C^/M^C^ and AF2 model reveal increased torsional twist in SARS-CoV-2 compared to SARS-CoV. (A) Cylinder and slabs diagram of SARS2-N/M (PDB ID 8ufm/A, this study) with backbone in light blue, helices in teal, and beta-strands in dark blue. Domains are labeled according to the complete structure of SARS1-N/M presented in Figure 2. Disulfide bond (C-C) between SARS2-N/M L516C-Y647C is indicated with the bond shown in yellow. Missing structure is shown as dashed lines. (B) Overlay of SARS2-N^C^M^C^ from 8ufm (light blue) and 8hbl (pink) is shown. Ellipse indicates structure 17N1 present in 8hbl. (C-E) Alignment of SUD-N domains of SARS2-N^C^M^C^ (light blue) with (C) SARS1-N/M 2w2g (pale yellow), (D) 8ufl (grey), and AF2 model of native SARS2-N/M (tan). Distance between disulfide bonds are indicated at arrows. Rotational twist is indicated in arrow, as measured from cysteine bond at center to surface exposed Lys587 that participates in binding to G4 quadruplex. In B-E, helix αM5 for each model is shown as tube to emphasize the relative orientation of the SUD-M domains (8ufm, teal; 2w2g, yellow; 8ufl, coral; AF2 brown) and rotation of panel images relative to each other. (E-H) Structural focus on linked LoopN8 with αM5 via cysteine bond or model with Leu and Tyr showing sidechain clash modeled by independent alignment of N and M domains.

The new SARS2-N^C^/M^C^ structure contained an interdomain disulfide bond formed between modified residues L516C from the 17N1 structure within LoopN8 and Y647C from αM5, just as found in SARS1-N/M. In the course of completing this work, similar structures of SARS2-N^C^/M^C^ were published with resolution of 1.35 Å (PDB ID 8gqc) and 8hbl 1.58 Å (PDB ID 8hbl) [21]. These structures are nearly identical except 8hbl resolved the 17N1 structure within LoopN8 (Figure 3B).

Similar to the recent report, the alignment of the SUD-N and SUD-M domains between SARS1-N/M structure 2w2g is rotated in SARS2-N^C^/M^C^. However, even as the individual domains are closely aligned, the overall r.m.s.d. is 9.3 Å. The high variance in structural alignment is due to rotation of SARS2-M by 21° relative to the orientation of these two domains in the previous determined SARS1-N/M structures (Figure 3C). The rotation occurs due to movement of LoopN8 such that the disulfide bond is 6.4 Å shifted compared to SARS1-N/M structures. When compared to our new complete structure 8ufl for SARS2-N/M, the r.m.s.d. is even greater at 12.8 Å with the rotation measured as 49°, the shift of the disulfide bond as 7.7 Å (Figure 3D), and flex of LoopN8 to an alternative orientation (Figure 3F-G). Further, the G4 quadruplex binding strand including Lys565 (SARS1-M) or Lys589 (SARS2-M) is significantly rotated away by 22.5 Å compared to structure 2w2g and 31.4 Å compared to structure 8ufl. Similarly, when the SUD-M domains are aligned, the PAIP1M binding region that includes Cys417 in SARS2-N and Cys393 in SARS1-N are shifted by 15 Å compared to 2w2g and 23.6 Å compared to 8ufl.

An AlphaFold2 (AF2) computational model was generated using ColabFold [26] to predict the structure of SARS2-N/M in the absence of the disulfide bond (Figure 3E). This AF2 predicted structure is identical to the x-ray determined structure in both domains (r.m.s.d. = 0.667 Å for SARS2-N and 0.673 Å for SARS2-M), but the AF2 model predicts the domains are dramatically reoriented by 104° rotation as measured from the aligned position of Cys516 in 8ufm and 8hbl to Lys589 in each construct. Residue Tyr516 in the AF2 model rotated 24.2 Å away from the position of Y516C in the determined SARS2-N^C^/M^C^. When the LoopN8 and αM5 structures were independently overlaid on the 8ufm structure (Figure 3H), it is revealed that in the absence of the two cysteines in SARS2-N/M, the two residues would clash and thus SUD-M was rotated away in the computational model. These data support that the C-C bond introduction provided a single conformation for crystallization in an otherwise highly flexible protein and the natural protein likely adopts a structure without the SUD-N/SUD-M interface due to clashes of the interface residues.

### Phylogenetic analysis of SUD-N/M reveals cysteines also absent in Pangolin-SARS and related coronaviruses

We next considered the question of whether the disulfide bond was a sequence variation gained by SARS-CoV or lost by SARS-CoV-2. A phylogenetic tree of protein sequences was generated using 14 diverse sequences of beta-coronaviruses available from the National Center for Biotechnology Information (NCBI). We found that the sequence sorted to three subclades. Of significance, in the SARS-CoV subclade (red in Figure 4), all of the sequences have both cysteines, including isolates from bats as well as from civets, the species thought to be the intermediate host for SARS-CoV. Moving along the phylogenetic tree of the SARS-CoV-2 subclade (blue in Figure 4), the cysteine in αM5 is changed to Tyr in a coronavirus isolated from a pangolin, the presumptive intermediate host for SARS-CoV-2. This strain has the cysteine in LoopN8, but the neighboring residue is changed from Ala to the bulkier Val that would likely also clash with the protruding Tyr from αM5. Both residues of LoopN8 then are changed to Asn in a more closely related pangolin isolate with the amino acid pair in a Bat SARS-like isolate and Bat RaTG13 having the Leu-Asn pair as found in SARS-CoV-2. Thus, the pair of residues in LoopN8 have varied as SARS-CoV-2 evolved in pangolins and bats resulting in loss of both cysteines.

**FIGURE 4.**
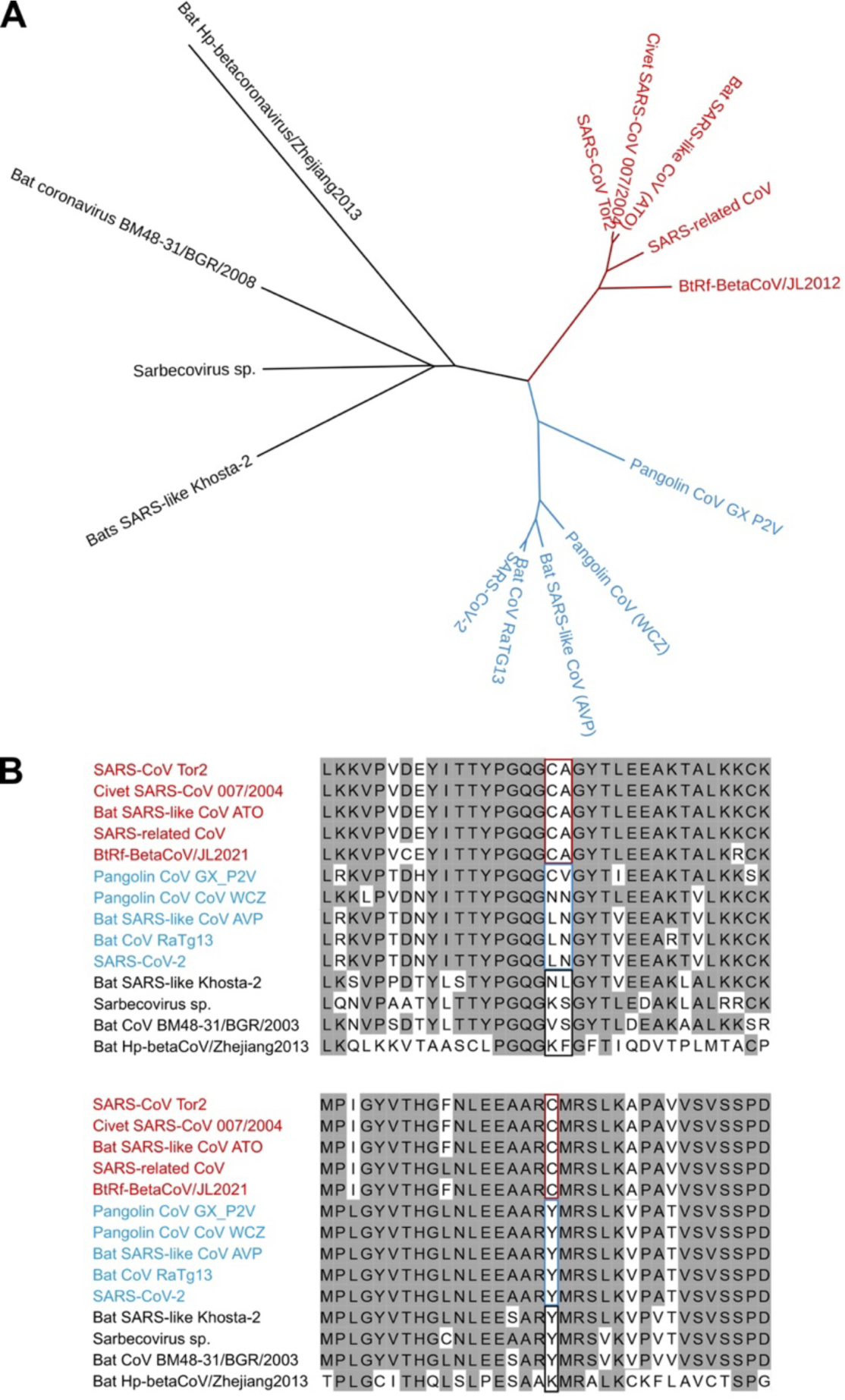
Phylogenetic relatedness of SUD-N/M protein sequences. (A) Unrooted phylogenetic tree showing relatedness of SUD-N/M protein sequences for 14 SARS-like betacoranaviruses with SARS-CoV subclade in red, SARS-CoV-2 subclade in blue, and other more distant SARS-related coronaviruses in black. (B) Amino acid sequences spanning the cysteine residues present in SARS-CoV structure as forming a double bond with text labels at left and square outlines colored as in panel A.

The third subclade is comprised of viral sequences more distantly related from the SARS-CoV and SARS-CoV-2 subclades (black in Figure 4). Interestingly, these also have the bulkier Tyr in αM5 and a variety of residues with bulky side chains extending from LoopN8. These data support that the C-C bond may be limited to the SARS-CoV subclade with bulky residues that would push apart SUD-N and SUD-M as the more common structure.

### The loss of the cysteine interaction does not impact function of SARS-N/M

A functional significance of the reorientation of SUD-N and SUD-M and the clash between residues Leu and Tyr that would occur if SARS2-N/M adopted the same structure as crystallized is that any contribution of SUD-N to G4 quadruplex binding would be lost. We confirmed that native SARS2-N^C^/M^C^ does bind *BCL2* G4 quadruplexes by zone interference gel electrophoresis with a K_d_ of 1.9 µM (Figure 5A). The binding efficiency was slightly improved compared to unmodified SARS2-N/M at 4.1 µM and essentially identical to K_d_ for SARS1-N/M at 1.6 µM. Further, this protein bound to SARS-CoV-2 5’-UTR at 5.3 µM, which is lower than the reported value for the unmodified protein at 41.4 µM in an experiment that was conducted simultaneously but split for purposes of publication [23]. These data support that the reintroduction of the cysteines may have slightly improved the binding affinity of the proteins for both G4 and for 5’-UTR with SUD-N.

**FIGURE 5.**
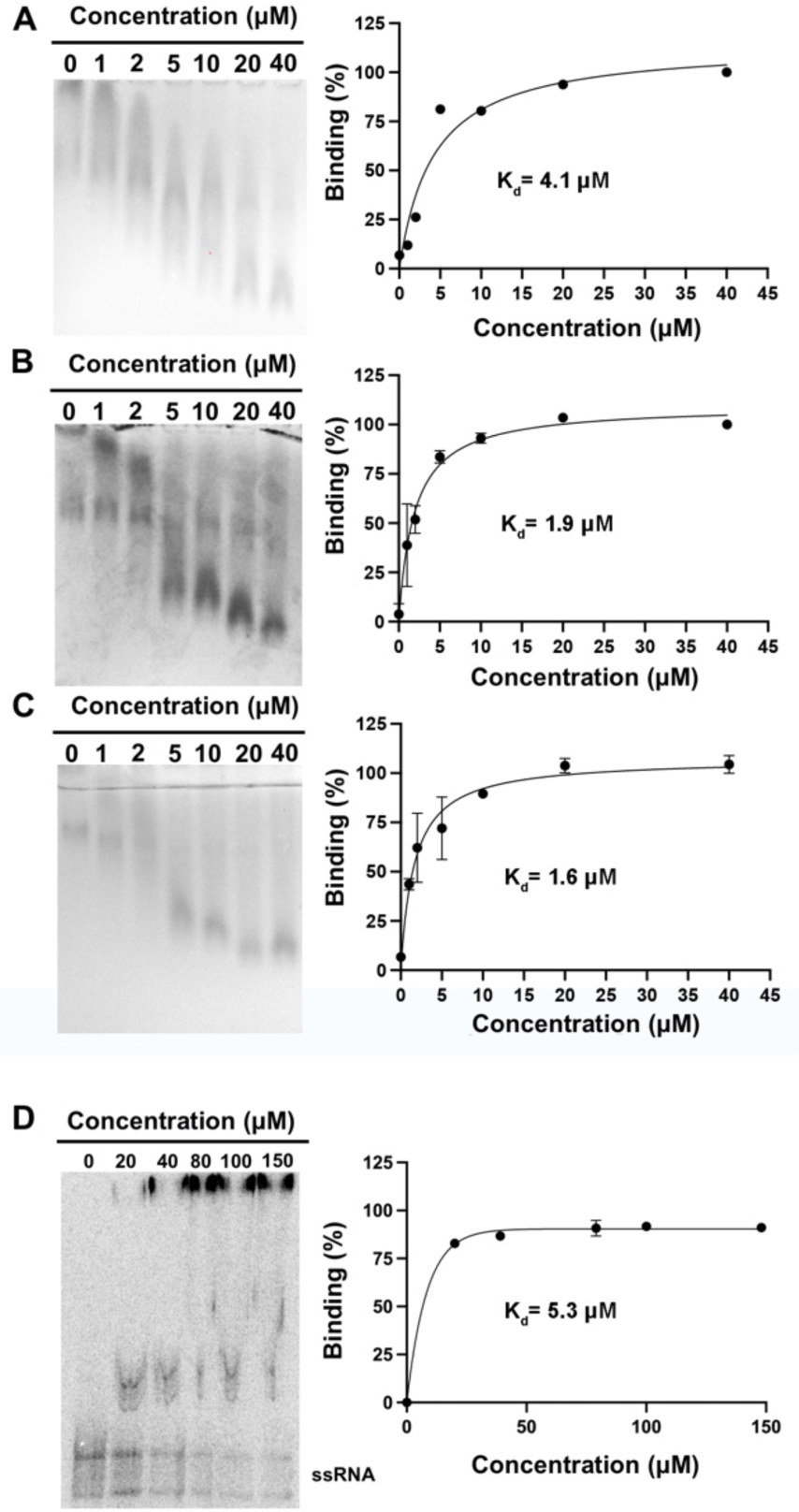
Introduction of Cys residues does not impact G4 quadruplex binding. (A-C) Zone-interference assay with protein as indicated mixed with Bcl-2 oligonucleotide folded into a G4 quadruplex the presence of 100 mM KCl in the presence of 0-40 µM proteins (A) SARS2-N/M, (B) SARS2-N^C^/M^C^, or (C) SARS1-N/M. (D) EMSA for quantification of binding of SARS2-N^C^/M^C^ to 5’-[^32^P]-labelled ssRNA of the SARS-CoV-2 5’-UTR (245 nt) region. For all panels, binding percentage (shown as a mean ±SD) and estimated K_d_ value were determined using three independent experiments. Representative gel shown at left and percent binding was plotted from triplicate assays at right.

As this domain is thought to also impact translational efficiency, we considered if the slight loss of nucleic acid affinity would impact viral replication. To test this, we used a SARS-CoV-2 replicon assay that uses luciferase as a readout of viral RNA replication in cells infected with single-round particles [27, 28]. The rationale for this experiment is that if translation was negatively impacted, protein complex for RNA replication would be reduced such that the overall copies of viral mRNAs would also be reduced. To test the impact of cysteines, the residues for both L516C and/or Y647C were introduced to the sequence of SARS-CoV-2 WA1 isolate. Although the mutations slightly reduced luciferase levels indicative of reduced viral replication when the cysteines were present, the reduction in luciferase signal was not significant in all three independently tested cells lines (Table 1). We further tested another mutation that could have introduced flexibility in the linker region. SARS-CoV has a proline at the position of SARS-CoV-2 Ser543 (Table 1). However, a S543P mutation either alone or in combination with L516C and Y647C also did not impact viral RNA replication in all three cell lines.

**Table 1.**
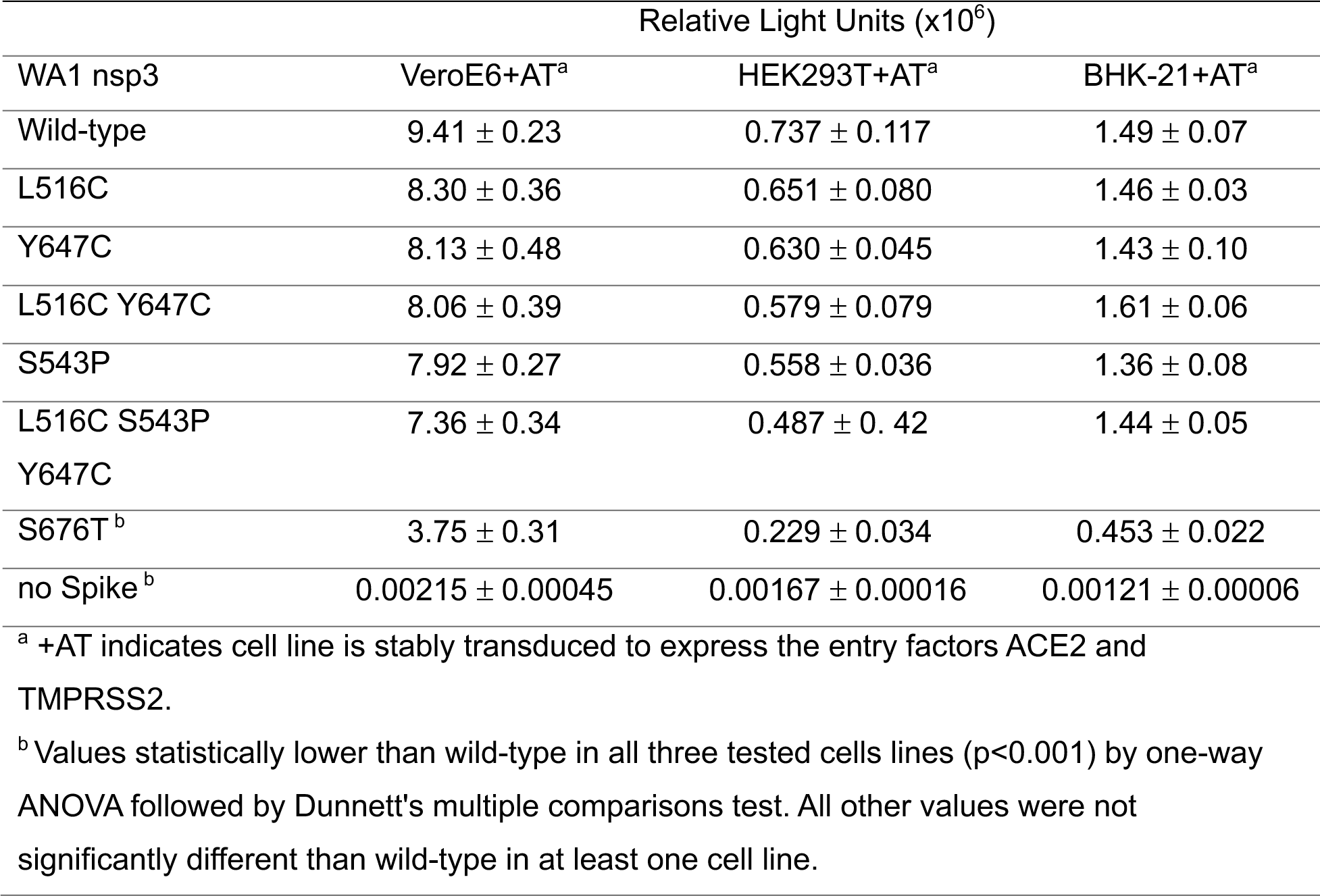
Impact of nsp3 SUD-N/M mutations on SARS-CoV-2 viral RNA replication.

As a control in the replicon assay, we tested a mutation of the last SUD-M residue Ser676 to threonine at the junction between SUD-M and DPUP (also known as SUD-C) that was previously reported to attenuate viral RNA replication [29]. We confirmed that the S676T mutation reduced viral replication by 60-70%. A Ser at this position is found in all 14 of the sarbecoviruses shown in Figure 4 and this residue is under negative selection in SARS-CoV-2 [30, 31], suggesting the residue is important for efficient coronavirus replication.

Altogether, these data support that SUD-N/M function as independent domains during viral mRNA replication. This increased flexibility may slightly reduce known activities of the unique domains in nucleic acid binding but thus did not confer a positive selective advantage based on the known activities of the unique domains in the emergence of SARS-CoV-2 as a human pathogen.

## DISCUSSION

Extensive studies of crystallization of SARS-CoV SUD-N/M have driven to development of hypotheses and conclusions that these two domains are interlinked by a disulfide bond that introduces rigidity. Docking studies originally supported that the interface between the two domains may even jointly contribute to binding to G4 quadruplexes, although mutagenesis studies support binding is related to a large loop adjacent to αM3 of only SUD-M [16, 20]. Our structures determined here, particularly a SARS-CoV structure in an altered conformation compared to all prior structures, reveals that there is torsional twist in the orientation of the two domains driven by flexibility of LoopN8 and the interdomain linker. The implication of these studies is that SUD-N/M were previously considered a “core” of the three unique domains, but our studies support that SUD-N and SUD-M are independent as is SUD-C.

Indeed, it is questionable if even in SARS-CoV and related SARS-like coronaviruses that also have both cysteines if disulfide bridges would occur within nsp3 in viral infected cells. Disulfide bonds do not form in the reducing environment of the cell cytosol into which nsp3 extends when if forms the crown of pore in the DMV. This leads to speculation of whether this disulfide bond is an in vitro artifact of overexpression and purification from *E. coli*. However, nsp3 does insert into endoplasmic reticulum membranes that ultimately form the DMV and thus these residues may access endoplasmic reticulum (ER) enzymes that introduce disulfide bonds at some point during DMV maturation [6]. Indeed recent data indicated the presence of antioxidant proteins present in viral replication vesicles isolated from Zika virus [32]. Since these vesicles likewise have an ER origin, a more reduced environment might modify nsp3 within the cell. Hence, for SARS-CoV this may be a functional stabilization during early establishment of the DMV but the bond may be broken in the cytoplasmic exposed crown. This stabilization however would not occur for SARS-CoV-2 nsp3.

A practical application of the introduction of the disulfide bond into SARS-CoV-2 SUD-N/M is that it may prove advantageous for advanced studies of SARS-CoV-2 nsp3 structure. Compilation of structural biology efforts have resulted in a near full coverage of SARS-CoV-2 nsp3 with x-ray structures of the independent domains. Recent work has described using cryoelectron tomography for visualization of the DMV crown with a resolution of 20 Å into which structures of Mac2 and Mac3 can be fitted [9, 10]. Notably, the structures fit to a bend in the crown structure. The lack of a C-C bond could introduce flexibility into this junction that could result into motion of the knob of each crown point, which is comprised of Ubl2 and Mac1 domains. The introduction of a C-C bond into nsp3 might be sufficient to stabilize nsp3 for the purpose of improving subatomic averaging to improve resolution presenting an advantage for more advanced structural studies. Indeed, introduction of the disulfide bond into an extended nsp3 comprising the entire N-terminal domain from Ubl1 to the first transmembrane domain did result in a more stable recombinant protein that was more easily purified than the unmodified protein [33]. This preliminary finding supports that flexibility of nsp3 may hamper advanced structural and biophysical studies of the large protein that could be improved by reintroduction of this bond found in some but not all SARS-like coronaviruses.

## METHODS

### Protein expression and purification

The primary amino acid sequence of nsp3 corresponding to SUD2-N/M and the SUD2-N^C^/M^C^ were taken from the original WA1 sequence 1231-1494 of for polyprotein 1ab, which is equivalent to the 413-678 of nsp3 from SARS-CoV-2 and residues:1231-1494 (polyprotein 1ab) equivalent to the SUD-N/M 389-652 of nsp3 and 1207-1470 of the polyprotein from SARS-CoV.

Nucleotide sequences encoding proteins were codon-optimized, using GenSmart™ free software (GenScript), synthesized and cloned into the pMCSG53 vector (Twist Biosciences), and transformed in *Escherichia coli* BL-21(DE3) Magic cells. The transformed bacteria were cultured in Terrific Broth media (SUD-N/M) or Se-Met (SUD2-N^C^/M^C^), and the protein expression was induced by addition of 0.5 mM isopropyl β-D-1-thiogalactopyranoside when cultures reached an optical density of ∼1.6-1.8 determined by the changes in the absorbance at 600 nm. The cells were collected by centrifugation at 5500 *xg* for 10 min, the pellets were resuspended in lysis buffer and frozen at -30 C until purification.

For protein purification, the frozen cells were thawed and sonicated at 45% intensity for 20 min, then treated with benzonase for 1 hr at 4 °C. The lysate was clarified by centrifugation at 30,000 *xg* for 40 min and loaded into a His-Trap FF [Ni–nitrilotriacetic acid (NTA)] column using a GE Healthcare ÅKTA Pure system using loading buffer (10 mM Tris-HCl (pH 8.3), 500 mM NaCl). The column was washed with loading buffer, followed by 10 mM Tris-HCl (pH 8.3), 500 mM NaCl, and 25 mM imidazole, and the protein was eluted with 10 mM tris (pH 8.3), 500 mM NaCl, and 500 mM imidazole. The protein was loaded onto a Superdex 200 26/600 column, ran with loading buffer, collected, and incubated with tobacco etch mosaic virus (TEV) protease overnight. The cleaved tag and TEV protease were separated from protein by Ni-NTA affinity chromatography using loading buffer and the protein was collected in the flow through. The protein was concentrated to 6.5 or 8.5 mg/ml and set up for crystallization immediately.

### Crystallization

The proteins in 0.3 M NaCl, 0.1 M Tris pH 8.3 (6.4 mg/ml for SARS1-N/M and 6.45 mg/ml for SARS2-N^c^/M^c^) were set up for crystallization as 2 µl crystallization drops (1-µl protein:1-µl reservoir solution) in 96-well (Corning) plates using commercially available Classics II, PEG’s II, AmSO_4_, Anions, and ComPAS Suites (Qiagen). Diffraction quality crystals were obtained for SARS1-N/M in Classics II screen (F6), 0.2 M ammonium sulphate, 0.1 M Bis-Tris pH 5.5, 25% PEG 3350 and for SARS2-N^c^/M^c^ in AmSO_4_ screen (A2), 0.2 M ammonium acetate, 2.2 M Ammonium sulfate. The crystals were cryoprotected in 2.0 M Lithium sulfate, and flash-frozen in liquid nitrogen for data collection.

### Data collection and refinement

Diffraction data were collected at the Life Science Collaborative Access Team (LS-CAT) at the Advanced Photon Source, Argonne National Laboratory. Data collection and refinement statistics are reported in Table 2. The data set was processed and scaled with the HKL-3000 suite [34]. The structure was solved by molecular replacement with Phaser [35] from the CCP4 suite using the crystal structure for SARS-CoV (PDB accession code 2w2g) as a search model. The residues of the linker region were removed and the model was split into two rigid bodies (domains) to use multiple body search algorithm in PHASER. The initial solution went through several rounds of refinement in REFMAC v5.8.0258 [36], and manual model corrections using Coot [37]. Well defined residues of the linker region were rebuilt and the models were further refined in REFMAC. The water molecules were generated using ARP/wARP [38], and ligands were added to the model manually during visual inspection in Coot. Translation-Libration-Screw (TLS) groups were created by the TLSMD server [39] (http://skuld.bmsc.washington.edu/~tlsmd/), and TLS corrections were applied during the final stages of refinement. Molprobity [40] (http://molprobity.biochem.duke.edu/) was used for monitoring the quality of the model during refinement and for the final validation of the structure. The final models and diffraction data were deposited to the Protein Data Bank (https://www.rcsb.org/) with the assigned PDB accession codes 8ufl and 8ufm.

**Table 2.** Structure determination data refinement.

| **Data processing** | **PDB ID 8ufl** | **PDB ID 8ufm** |
| --- | --- | --- |
| Structure | SARS1-N/M | SARS1-N<sup>c</sup>/M<sup>c</sup> |
| Beamline | APS 21-ID-F | APS 21-ID-D |
| Wavelength (Å) | 0.97872 | 1.12704 |
| Resolution range (Å) | 30.00-2.50 | 30.00-1.65 |
| Space group | *P*2<sub>1</sub>2<sub>1</sub>2<sub>1</sub> | *P*3<sub>1</sub>21 |
| Cell parameters (Å, °) | *a*=67.70, *b*=84.86, *c*=93.60;<br>$\alpha$=90.00; $\beta$=90.00; $\gamma$=90.00 | *a*=86.27, *b*=86.27, *c*=76.67;<br>$\alpha$=90.00; $\beta$=90.00; $\gamma$=120.00 |
| Unique reflections | 19,083 (908) | 40,007 (1,971) |
| Multiplicity | 5.6 (5.6) | 18.8 (14.4) |
| Completeness (%) | 100.0 (100.0) | 99.9 (99.7) |
| Mean I/sigma(I) | 15.1 (2.2) | 30.1 (2.0) |
| Wilson B-factor (Å<sup>2</sup>) | 41.1 | 26.8 |
| R-merge<sup>a</sup> | 0.112 (0.800) | 0.100 (1.625) |
| R<sub>pim</sub> | 0.052 (0.370) | 0.024 (0.437) |
| CC1/2<sup>b</sup> | 0.97 (0.74) | 1.00 (0.77) |
| **Refinement** |  |  |
| Resolution range (Å) | 29.80-2.51 (2.57-2.51) | 28.26-1.65 (1.69-1.65) |
| Reflections work/test | 18,109(1,262)/928(70) | 37,981(2,743)/1,997(153) |
| R<sub>work</sub>/R<sub>free</sub><sup>c</sup> | 0.214(0.307)/0.267(0.314) | 0.185(0.324)/0.208(0.342) |
| Total number of atoms | 4,268 | 2,209 |
| macromolecule atoms | 4,044 | 1,921 |
| Ligand/solvent (H<sub>2</sub>O) | 66/125 | 42/208 |
| RMSD (bonds) (Å) | 0.004 | 0.006 |
| RMSD (angles) (°) | 1.369 | 1.454 |
| Ramachandran favored (%) | 95.0 | 98.0 |
| Ramachandran allowed (%) | 5.0 | 2.0 |
| Ramachandran outliers (%) | 0.0 | 0.0 |
| Rotamer outliers (%) | 0.2 | 0.0 |
| Clashscore | 6 | 1 |
| Average B-factor (Å<sup>2</sup>) | 48.2 | 37.2 |

### Sequence and Structure analysis

All structure analysis was conducted using UCSF ChimeraX software including Matchmaker, Distances, and Structure Alignment tools [41]. AF2 models were generated in ChimeraX using its interface to ColabFold [26]. Sequence alignment and phylogeny were conducted in MacVector v.18.2.5 using sequences sourced from the National Center for Biotechnology Information (ncbi.nlm.nih.gov) with a tree figure produced using iTOL v.6 (itol.embl.de). NCBI accession number for sequences used were for comparison were: SARS-CoV Tor2 AFR58699.1, Civet SARS-CoV 007/2004 AAU04645.1, Bat SARS-like CoV ATO98203.1, SARS-related CoV WCZ55791.1, BtRf-BetaCoV/JL2012 AIA62276.1, Pangolin CoV GX_P2V QVT76604.1, Pangolin CoV WCZ55784.1, Bat SARS-like CoV AVP78030.1, Bat CoV RaTG13 QHR63299.2, Bats SARS-like Khosta-2 QVN46568.1, Sarbecovirus sp. QTJ30152.1, Bat coronavirus BM48-31/BGR/2008, and Bat Hp-betacoronavirus/Zhejiang2013.

### Zone-interference gel electrophoresis (ZIGE)

To measure the interaction of SUD-N/M from SARS-CoV and SARS-CoV-2 a zone-interference gel electrophoresis was performed, using a custom 3D-printed comb, similar to the reported by Abrahams et al [42]. and Tan et al [12]. Briefly, 10 µM of each protein was incubated with 0-40 µM of the folded *BCL2* promotor in the presence of 100 mM KCl at room temperature for 30 min. The agarose gel (1%) gel was loaded with 0-40 µM of the ligand in TBE (20 mM Tris, 50 mM boric acid, 0.1 mM ethylendiaminotetracetic acid (EDTA), pH 8.3), 1% dimethylsulfoxide (DMS0) and 0.01% of bromophenol blue (BPB) in the long slots and run at 100 mA for one minute or until the front of the dye reached the small slot. Then the samples of protein with ligand were mixed with 1% DMSO and 0.01% BPB, loaded in the small slots, and continued the electrophoresis in TBE for 1 hr. The gel was fixed using 40% acetic acid 30% for 2 hr ethanol and stained with 0.2% Coomassie blue in for 1h and distained with until background washed out, and imaged. The K_d_ was calculated by measuring the protein migration from the small well to the front of the agarose gel, using 100% as the migrated distance at maximum concentration (40 µM). The K_d_ was determined by non-linear fit, one binding using Prism (GraphPad V10).

### Electrophoretic mobility shift assays (EMSA)

The reaction mixture for RNA binding assays contained 50 mM Tris-HCl (pH 8.0), 150 mM NaCl, 5 mM CaCl_2_, 1 mM dithiolthreitol (DTT), 20 U RNaseOUT (Invitrogen), and 8 nM 5’-[^32^P]-labelled RNA substrate. Reactions were incubated for one hour at 37°C, quenched by the addition of glycerol loading dye, and separated on 6% native polyacrylamide gels. Results were visualized using a Phosphoimager, with the percentage of bound substrate quantified using ImageLab software (Bio-Rad). Values were plotted against total protein concentration to determine K_d_ values using non-linear regression fit in Prism software (GraphPad).

### SARS-CoV-2 Replicon Assay

The SARS-CoV-2 replicon assay was conducted as described previously [27]. Briefly, the pBAC SARS-CoV-2 ΔSpike WT or nsp3 amino acid modified plasmids (1 μg), were transfected into BHK-21 cells along with N R203M and S Delta variant expression vectors (0.5 μg each) in a 24-well tissue culture dish. The culture media was replaced with fresh growth medium 12–16 hours post-transfection. The media containing single-round infectious particles was collected and 0.45 μm-filtered 72 hours post-transfection. The filtered media containing single-round infectious particles was added to 2x10^4^ VeroE6 cells stably expressing ACE2 and TMPRSS2 (Vero+AT), HEK293T cells stably expressing ACE2 and TMPRSS2 (293T+AT), and BHK-21 cells stably expressing ACE2 and TMPRSS2 (BHK21+AT) cells in a 96-well plate. The cells were washed with 200 µL culture medium and 100 µL culture medium was added 12–24 hours post-infection. To measure luciferase activity, 50 µL of superrnatant from infected cells was mixed with an equal volume of Nano-Glo luciferase assay buffer and substrate and analyzed on an Infinite M Plex plate reader (Tecan).

## SUPPLEMENTARY MATERIAL DESCRIPTION

There are no supplemental materials.

## DATA AVAILABILITY

Atomic coordinates and related experimental data were deposited and released to the Worldwide Protein Data Bank (wwPDB) at rcsb.org with accession numbers 8ufl and 8ufm. Plasmids for expression of SUD1-N/M, SUD2-N/M, and SUD2-N^C^/M^C^ are available from Karla Satchell upon request.

## AUTHOR CONTRIBUTIONS

MRL designed the project and conducted experiments. GM and JB solved the crystal structures. TYT, SL, SY, LS, JR, and KK conducted experiments. HSS consulted on project development. AS, PJS, MO, and KJFS supervised research and provided funding. KJFS prepared final manuscript for publication with input from MRL, GM, and TYT.

## ACKNOWLEDGEMENTS

We thank A. Creanga and B. Graham for the Vero cells overexpressing human ACE2 and TMPRSS2. This project was supported by HHS/NIH/NIAID contract 75N93022C00035 (to K.J.F.S. and A.S.) and funding from the University of Toronto COVID-19 Action Initiative (to P.S.). Project further supported by NIH grants R37 AI033493 and R01 AI146073 (to H.S.) and Roddenberry Foundation, P. and E. Taft and the Pendleton Foundation (to M.O). M.O. is a Chan Zuckerberg Biohub – San Francisco Investigator. M.R.L. is a mentored Principal Investigator at the Autophagy, Inflammation and Metabolism Center of Biomedical Research Excellence (NIH: P20GM121176). The research used resources of the Advanced Photon Source, a U.S. Department of Energy (DOE) Office of Science User Facility operated for the DOE Office of Science by Argonne National Laboratory under Contract No. DE-AC02-06CH11357. Use of the LS-CAT Sector 21 was supported by the Michigan Economic Development Corporation and the Michigan Technology Tri-Corridor (Grant 085P1000817).

